# Transcranial stimulation of alpha oscillations upregulates the default mode network

**DOI:** 10.1101/2021.06.11.447939

**Authors:** Kevin J. Clancy, Jeremy A. Andrzejewski, Jens T. Rosenberg, Mingzhou Ding, Wen Li

## Abstract

The default mode network (DMN) is the most prominent intrinsic connectivity network, serving as a key architecture of the brain’s functional organization. Conversely, dysregulated DMN is characteristic of major neuropsychiatric disorders. However, the field still lacks mechanistic insights into the regulation of the DMN and effective interventions for DMN dysregulation. The current study approached this problem by manipulating neural synchrony, particularly, alpha (8-12 Hz) oscillations, a dominant intrinsic oscillatory activity that has been increasingly associated with the DMN in both function and physiology. Using high-definition (HD) alpha-frequency transcranial alternating current stimulation (*α*-tACS) to stimulate the cortical source of alpha oscillations, in combination with simultaneous EEG-fMRI, we demonstrated that *α*-tACS (vs. sham control) not only augmented EEG alpha oscillations but also strengthened fMRI and (source-level) alpha connectivity within the core of the DMN. Importantly, increase in alpha oscillations mediated the DMN connectivity enhancement. These findings thus identify a mechanistic link between alpha oscillations and DMN functioning. That transcranial alpha modulation can upregulate the DMN further highlights an effective non-invasive intervention to normalize DMN functioning in various disorders.

**Significance Statement:** In the brain’s functional organization, the default mode network (DMN) represents a key architecture, whose dysregulation is involved in a host of major neuropsychiatric disorders. However, insights into the regulation of the DMN remain scarce. Through neural synchrony, the alpha-frequency oscillation represents another key underpinning of the brain’s organization and is thought to share an inherent interdependence with the DMN. Here, we demonstrated that transcranial alternating current stimulation of alpha oscillations (*α*-tACS) not only augmented alpha activity but also strengthened connectivity of the DMN, with the former serving as a mediator of the latter. These findings reveal that alpha oscillations can support DMN functioning. In addition, they identify an effective non-invasive approach to regulate the DMN via *α*-tACS.

## Introduction

It is widely recognized that the brain self-organizes into large-scale intrinsic networks. Such intrinsic organization is so fundamental to normal neural functioning that it commands 60-80% of the brain’s energy (1). Two main mechanisms—intrinsic inter-regional connectivity and inter-neuronal synchrony—are thought to underpin the brain’s organization (2-6). The default mode network (DMN), emerging from intrinsic inter-regional connectivity crisscrossing a large extent of the brain, occupies the apex of intrinsic connectivity networks (7) and dominates the brain’s intrinsic activity (1, 8). Accordingly, the DMN supports advanced human mental faculties (e.g., consciousness, self-reference, social inference, remembering the past and expecting the future) (8) while its dysregulation is characteristic of major neuropsychiatric disorders (e.g., Alzheimer’s disease, schizophrenia, posttraumatic stress disorder) (9-11). However, mechanisms regulating the DMN remain elusive while effective interventions for DMN dysregulation are lacking.

Inter-neuronal synchrony is thought to be inherently related to inter-regional connectivity and potentially bind and sculpt such connectivity through neural development (2-6). Importantly, the alpha (8-12 Hz) oscillation, the primary rhythm of intrinsic neural synchrony (12, 13), has been linked to the DMN functioning (4, 14). In fact, rapid advances in neuroimaging and neurocomputing have brought forward mounting evidence of multifaceted physiological and functional associations between the alpha oscillation and the DMN. Physiologically, resting-state (RS) simultaneous EEG-fMRI (electroencephalography and functional magnetic resonance imaging) recordings have revealed intrinsic positive coupling between alpha oscillations and DMN activity (15-19). Of particular relevance, alpha oscillations are found to be the primary neural synchrony linking the posterior and anterior hubs of the DMN (the posterior cingulate cortex/PCC and medial prefrontal cortex/mPFC, respectively) (14, 18, 20, 21). Functionally, alpha oscillations and the DMN are both involved in disengaging the brain from the sensory environment and maintaining the resting state (1, 12, 22) while alpha desynchrony and DMN dysconnectivity, including specific disruption of alpha-oscillatory PCC-mPFC connectivity, often co-occur in major neuropsychiatric disorders (23-25).

Nonetheless, these associations between alpha oscillations and the DMN remain correlational in nature, calling for experimental investigation to ascertain their mechanistic linkage. Owing to the proximity of its primary source (the occipitoparietal cortex) to the scalp, the alpha oscillation is highly responsive to transcranial stimulation (26-29) and can be a viable target for experimental manipulation. Among the many transcranial stimulation technologies, transcranial alternating current stimulation (tACS) applies frequency-specific sinusoidal electric currents through the scalp, which is uniquely advantageous in mimicking and entraining endogenous oscillations by tuning not only the frequency and amplitude but also the oscillatory phase. The latter, by enhancing phase synchronization, could be particularly effective at facilitating inter-regional connectivity (30). Therefore, we experimentally manipulated alpha oscillations with MR-compatible high-definition (HD) alpha-frequency tACS (*α*-tACS) targeting the occipitoparietal alpha source. Contrasting simultaneous EEG-fMRI before and after tACS (vs. sham control), we sought to demonstrate DMN upregulation via the enhancement of alpha oscillations.

## Results

### Validation of tACS (*α-tACS increased alpha power and connectivity*)

We applied high-definition (HD) tACS using a 4 × 1 montage over midline occipitoparietal sites (with 4 surrounding + 1 central electrodes forming a closed circuit; Fig. 1*A*), which were selected to maximally target the primary alpha cortical source—the occipitoparietal cortex (12, 13, 27). Previous work has demonstrated the efficacy and reproducibility of this tACS protocol in augmenting both alpha local power and long-range connectivity (27). Finite-element model simulation of the current distribution based on a standard head model confirmed maximal electric fields (0.21 V/m) in the occipitoparietal cortex (peaking at the occipito-parietal junction; x, y, z = 16, -86, 36), relative to minimal electric fields (< 0.02 V/m) in frontal regions. Another finite-element model estimation based on a realistic head model (the average T1 of the Active group) yielded a similar distribution of the current (*SI Appendix* Fig. *S*1). Source-level analysis of EEG alpha power in the Active group showed maximal alpha power increase in the right occipitoparietal cortex (Fig. 1*B*). Importantly, in support of the efficacy of α-tACS, we observed increases in right posterior alpha power and right posterior-to-frontal alpha connectivity in the Active (vs. Sham) group from Pre-to Post-stimulation (Fig. 1*C-G*). These effects, including the right-hemisphere dominance, replicated our previous tACS findings (27).

**Figure 1.**
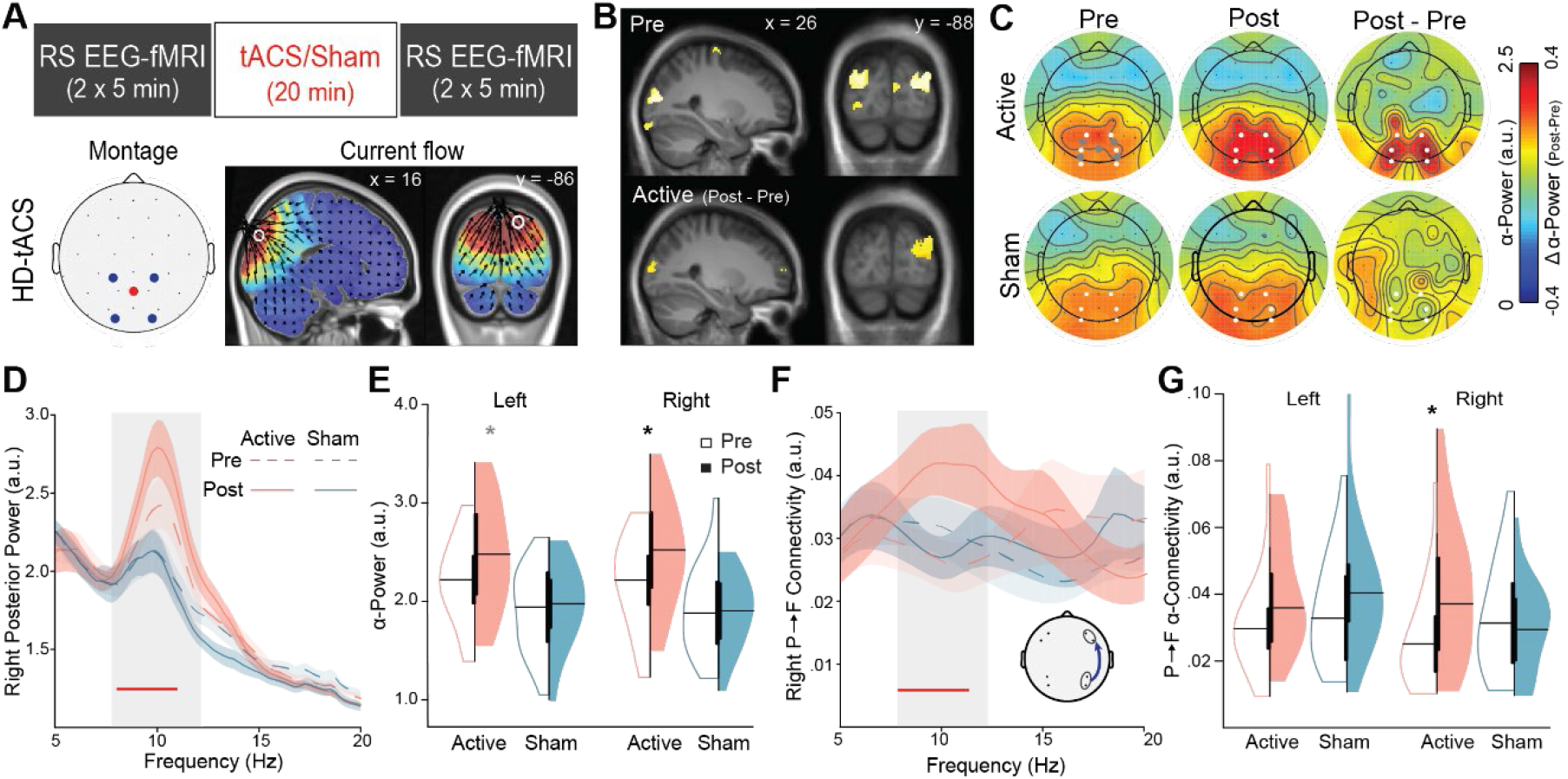
α-tACS efficacy. **A**) Experimental design. Participants underwent RS EEG-fMRI recordings immediately before and after tACS/Sham stimulation. High-definition (HD) tACS was administered over the occipitoparietal midline with a 4 ⨯ 1 montage (4 surrounding + 1 central electrodes forming a closed circuit). Finite-element modeling of the current flow based on a standard head model indicated maximal and focal electric fields (0.21 V/m) in the occipitoparietal cortex (peaking at the occipito-parietal junction, as indicated by the white circle; x, y, z = 16, -86, 36; Montreal Neurological Institute coordinates), relative to minimal electric fields (< 0.02 V/m) in frontal regions. **B)** We further localized the source of alpha power to the bilateral occipitoparietal cortex (display threshold *p* < 0.001 FWE, k = 20; Top row). Source-level alpha power increase after tACS was further localized to the right occipitoparietal cortex (display threshold *p* < .05, k = 10; Bottom row). **C**) Topography of alpha (8-12 Hz) power before and after stimulation. White dots indicate left and right occipitoparietal electrodes from which alpha power was extracted. **D**) Spectral waveforms averaged across the right occipitoparietal electrodes demonstrate specific increases in the Active (vs. Sham) group, which were restricted to the alpha frequency (grey box); the red bar indicates frequency bins (0.25 Hz each) showing significant tACS effects; Ribbon = SEM. **E**) Violin plots for alpha power over left and right electrodes indicate significant right posterior power increase in the Active group. **F**) Right hemisphere posterior→frontal (P→F) Granger causality (GC) waveforms demonstrate specific increases in the Active (vs. Sham) group, which were restricted to the alpha frequency (grey box); the red bar indicates frequency bins (0.5 Hz each) showing significant tACS effects; the inset shows ipsilateral electrode pairs used for alpha GC as used in our prior studies (27, 59, 60); Ribbon = SEM. **G**) Violin plots for alpha-frequency P→F GC in the left and right hemispheres indicate significant right hemispheric GC increase in the Active group. \**p < 0*.*05* (black/gray: surviving double/single contrast).

Specifically, after tACS, the Active group (*n* = 17) showed significant increase in right posterior alpha power from the baseline (*t* = 2.69, *p* = 0.015) while no change was observed in the Sham group (*n* = 19; *p* = 0.92; Fig. 1*C-E*). A double contrast (Post - Pre_Active – Sham_) further confirmed specific alpha power increase in the Active (vs. Sham) group (*t* = 2.14, *p* = 0.040). Left posterior alpha power also increased in the Active group (*t* = 2.16, *p* = 0.045), which, however, failed to survive the double contrast (*t* = 1.36, *p* = 0.184). As evinced by recent neural computational and electrophysiological (including intracranial recordings) studies, alpha projections track a selective posterior-to-frontal (P→F) direction (i.e., directed alpha P→F connectivity), via posterior-to-frontal cortical synchronization or travelling waves (14, 20, 21, 31-35). We thus used Granger causality (GC) analysis to examine changes in this alpha connectivity. We found that the Active group also exhibited an increase in right-hemispheric alpha P→F connectivity (*t* = 2.36, *p* = 0.031), which was again absent in the Sham group (*p* = 0.621; Fig. 1*F-G*). A similar double contrast confirmed that this alpha connectivity increase was specific to the Active group (*t* = 2.12, *p* = 0.042). Exploratory analyses of opposite F→P connectivity showed no effect of tACS (*SI Appendix*). Finally, as illustrated in Fig. 1*C&F*, both power and connectivity increases were constrained to the alpha frequency, highlighting the specific tACS effects on alpha oscillations.

### α-tACS increased DMN BOLD connectivity

Extracting RS fMRI BOLD timeseries from the DMN regions of interest (ROIs)—midline hubs (mPFC and the ventral and dorsal subdivisions of PCC—vPCC and dPCC) and a key lateral node (left/right angular gyrus/ANG)—before and after stimulation, we conducted ROI-based functional connectivity analysis (followed by multiple comparison correction based on the false discovery rate/FDR). The Active group demonstrated increases in vPCC-mPFC (*t* = 2.87, *p* = 0.011) and vPCC-rANG (*t* = 3.12, *p* = 0.007) connectivity after tACS (Fig. 2*A-C*). There was no change in the Sham group (*p*’s > 0.13) while double contrasts (Post - Pre_Active – Sham_) further confirmed these increases were specific to the Active group (*t*’s > 2.79, *p’s* < 0.009, FDR *p* < 0.05). Whole-brain vPCC-seed connectivity maps (of both simple and double contrasts) confirmed these results and, importantly, indicated that the increases were largely constrained to the DMN (Fig. 2*C*; *SI Appendix*; Fig. *S*2).

**Figure 2.**
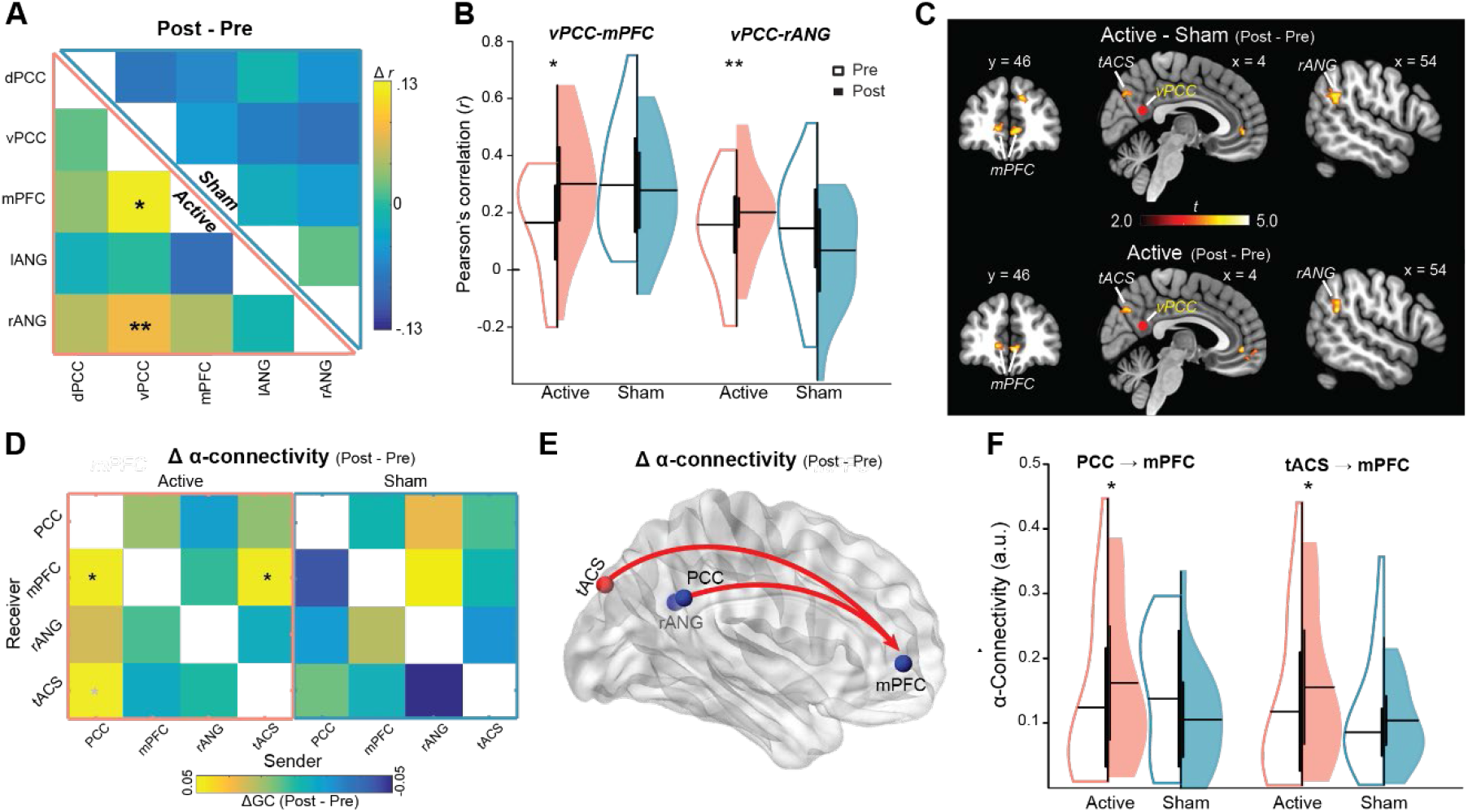
α-tACS enhanced DMN BOLD and alpha connectivity. **A**) Post-vs. pre-stimulation connectivity matrix for the Active (lower half) and Sham Control (upper half) groups and **B**) their respective violin plots demonstrate that tACS increased DMN connectivity (vPCC-mPFC and vPCC-rANG). **C)** vPCC-seed whole-brain connectivity maps further indicate that increases in vPCC connectivity (in both the double and single contrasts) were largely restricted to the DMN (bilateral mPFC, rANG) and the tACS site (display threshold at *p* < .005 uncorrected, *k* > 10). **D & E)** The Active group further demonstrated increases in alpha-frequency PCC→mPFC and tACS_site_→mPFC connectivity (GC), which also survived the double contrast (vs. changes in the Sham group). **F)** Their respective violin plots. \**p < 0*.*05, **p<0*.*01*, (black/gray: surviving double/single contrast); black/solid line = double contrast, gray /dotted line = single contrast. Red dot indicates vPCC seed.

### α-tACS increased DMN alpha-frequency connectivity

We then examined the effect of tACS on source-level (DMN) alpha P→F connectivity. A double contrast (Post - Pre_Active – Sham_) showed a specific increase in alpha PCC→mPFC connectivity in the Active (vs. Sham) group (*t* = 2.27, *p* = 0.030; Fig. 2*D-F*) after stimulation. Specifically, the Active group showed an increase (*t* = 1.74, *p* = 0.049, one-tailed) while the Sham group trended towards a decrease (*t* = -1.48, *p* = 0.079, one-tailed) in this connectivity. The alpha rANG→mPFC connectivity was not affected by tACS (*p* = 0.92). Exploratory analyses of alpha horizontal connectivity between rANG and PCC (in both directions) and opposite F→P alpha connectivity (i.e., mPFC→PCC and mPFC→rANG) showed no effect of tACS (*p*’s > 0.440).

### Alpha increase mediated DMN connectivity enhancement via α-tACS

Correlational analysis showed that increases in BOLD vPCC-mPFC connectivity after tACS strongly correlated with increases in alpha P→F connectivity (*r* = 0.59, *p* < 0.001; Fig. 3*A*). Changes in P→F connectivity in other (delta, theta, and beta) frequencies showed no such association (*p*’s > 0.111), demonstrating a unique association between changes in alpha-frequency connectivity and DMN connectivity. Furthermore, increases in BOLD vPCC-rANG did not correlate with alpha connectivity (*r* = 0.14, *p* = 0.545), highlighting the association of alpha P→F connectivity with DMN posterior-anterior connectivity. Importantly, a mediation analysis of tACS modulation of BOLD vPCC-mPFC connectivity revealed a significant indirect effect of alpha P→F connectivity (beta = 0.076, CI = [0.018 0.189]), suggesting that increases in alpha P→F connectivity mediated the effect of tACS on BOLD vPCC-mPFC connectivity (Fig. 3*B*).

**Figure 3.**
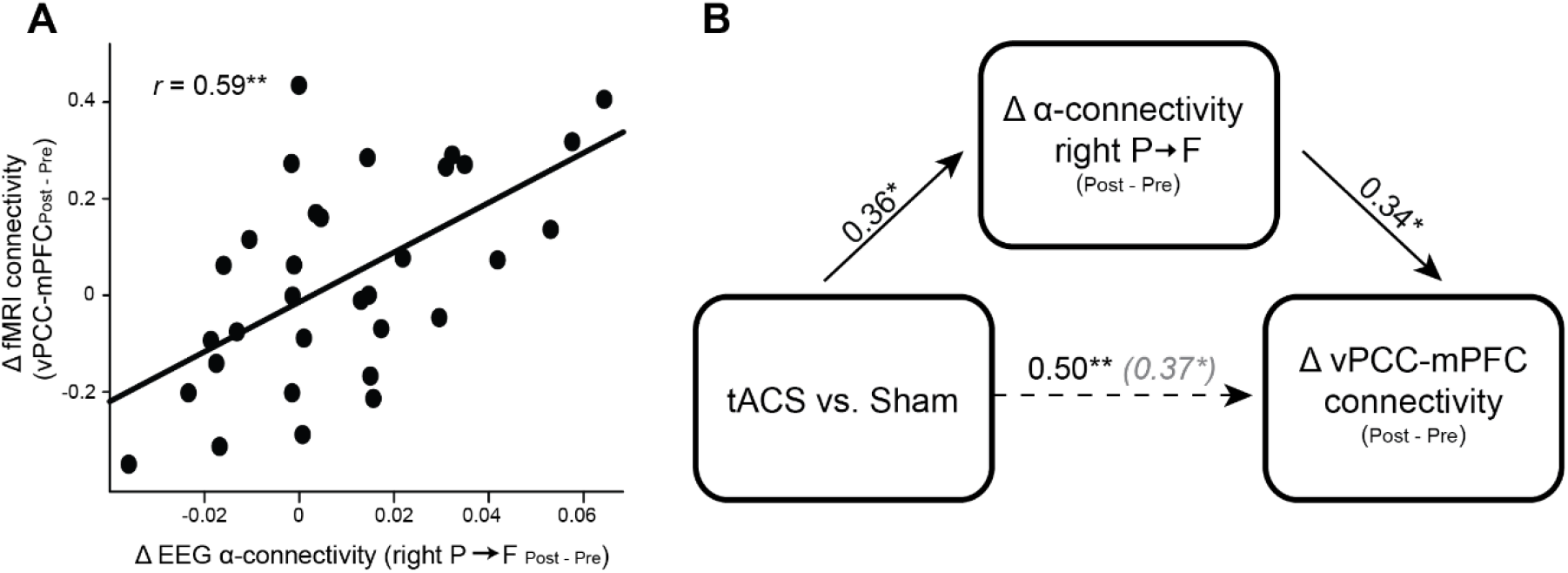
Alpha P→F connectivity increase mediated DMN connectivity enhancement via α-tACS. **A)** Scatterplot demonstrating a positive correlation between changes in BOLD vPCC-mPFC connectivity and right hemisphere P→F alpha-frequency connectivity/GC. **B)** Mediation model demonstrating indirect (i.e., mediation) effect of tACS group on increases in DMN connectivity through increases in right hemisphere P→F alpha-frequency GC. Path strengths are indicated by standardized beta coefficients. The parenthetical beta coefficient reflects the direct path strength after controlling for changes in P→F GC. \**p* < 0.05, \*\**p* < 0.005.

### α-tACS increased connectivity between the tACS site and the DMN

To attain mechanistic insights into tACS-induced neuromodulation, we examined connectivity between the tACS site (a 10-mm sphere around the voxel with maximal electrical field of tACS; Fig. 1*A*) and the DMN. Active group demonstrated vPCC-tACS_site_ connectivity increase after tACS (*t* = 3.73, *p* = 0.002) while the Sham group showed no change (*p* = 0.578; Fig. 4*A*). A double contrast further confirmed the specific increase in the Active (vs. Sham) group (*t* = 2.30, *p* = 0.028). The Active group showed tACS_site_-mPFC connectivity increase after tACS (*t* = 2.89, *p* = 0.011) while the Sham group showed no change (*p* = 0.836). Again, a double contrast confirmed the increase was specific to the Active (vs. Sham) group (*t* = 1.75, *p* = 0.045 one-tailed). No effects of tACS emerged in the tACS_site_-rANG connectivity (*p*’s > 0.272). Whole-brain tACS_site_ seed-based connectivity analysis confirmed these results while showing little connectivity change outside the DMN (*SI Appendix*; Fig. 4*B*).

**Figure 4.**
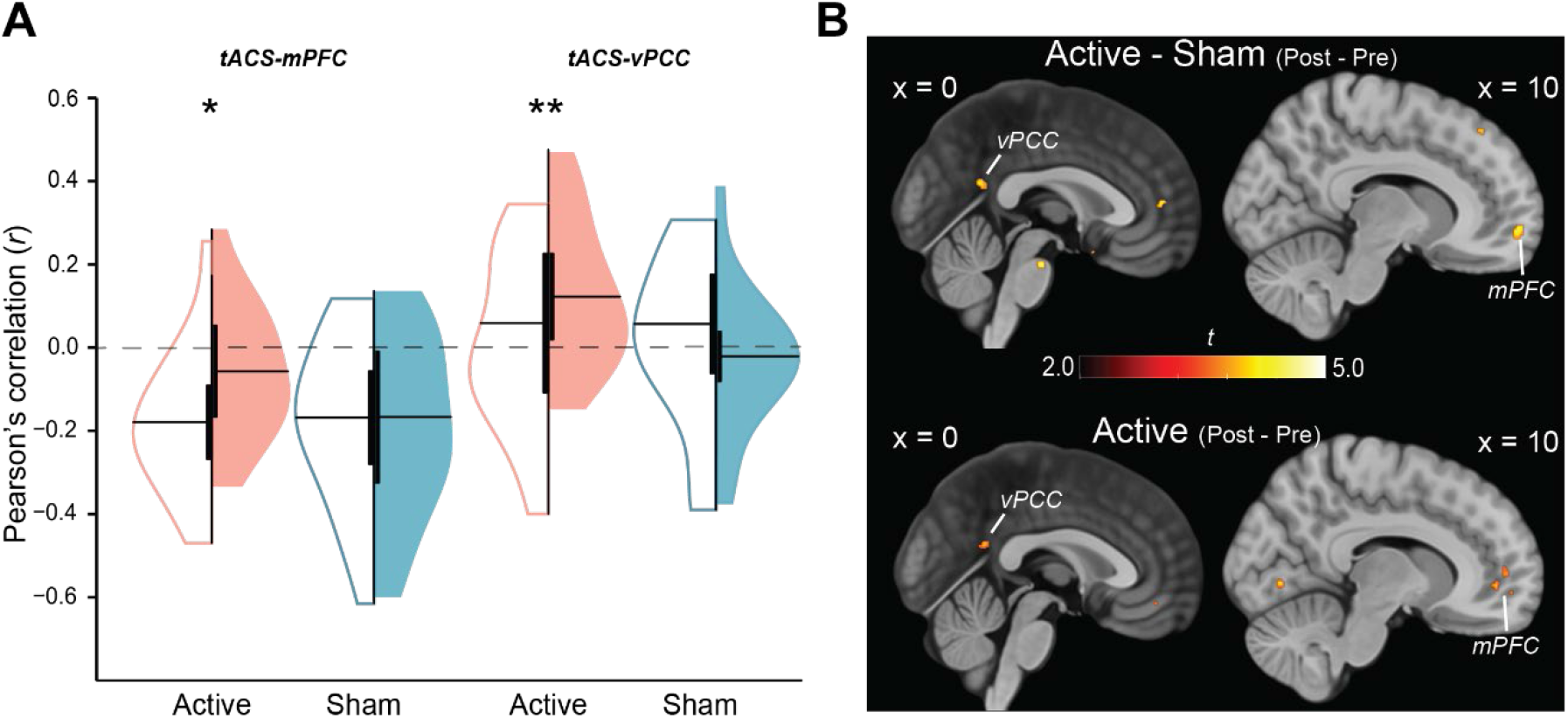
Increased connectivity between tACS site and DMN. **A)** Violin plots demonstrate that tACS increased connectivity between the tACS site and midline DMN hubs (tACS-mPFC and tACS-vPCC). **B)** tACS_site_ seed-based whole-brain connectivity maps further indicate increases in tACS connectivity to midline DMN hubs for the Active group alone (*p* < 0.005 uncorrected, *k* > 10). \**p* < 0.05, \*\**p* < 0.01.

We further examined alpha-frequency connectivity changes from the tACS site to the DMN. Source-level alpha causality analysis indicated increased tACS_site_→mPFC (but not tACS_site_→PCC or tACS_site_→rANG) alpha connectivity in the Active group (*t* = 1.89, *p* = 0.038, one-tailed; Fig. 2*D*), which a double contrast indicated to be specific to the Active (vs. Sham) group (*t* = 1.64, *p* = 0.056 one-tailed). Akin to the posterior-to-frontal alpha projection, these results suggest that α-tACS of the occipitoparietal cortex could drive alpha oscillations in the mPFC.

## Discussion

Combining MR-compatible HD α-tACS with simultaneous EEG-fMRI, we demonstrated that α-tACS of the alpha source in the occipitoparietal cortex not only augmented alpha oscillations but also strengthened BOLD and alpha-frequency oscillatory connectivity within the DMN. Importantly, tACS-induced augmentation of P→F alpha connectivity mediated the tACS enhancement of BOLD connectivity between DMN hubs (vPCC-mPFC). By contrast, no tACS effects emerged outside the alpha frequency or the DMN. These findings thus indicate that the DMN can be upregulated by transcranial stimulation of alpha oscillations. Moreover, they provide experimental evidence to mechanistically link alpha oscillations, particularly alpha connectivity, to the DMN, lending further credence to the notion of alpha-DMN interdependence (4, 14).

By strengthening oscillatory circuits via spike-timing dependent plasticity (STDP) and long-term potentiation (LTP) at the synapse (36, 37), α-tACS can have lasting effects on long-range BOLD (38-40) and alpha-frequency oscillatory connectivity (27, 29). However, network-level effects have been examined in only a handful α-tACS-fMRI studies, which, using varied stimulation montages and protocols, have yielded ambiguous findings (38-40). We chose to apply HD-tACS for concentrated stimulation of the alpha source in the occipitoparietal cortex, which was demonstrated to amplify not only the local alpha power but also the posterior-to-frontal alpha connectivity (27). As such, this tACS protocol would capitalize on the selective posterior-to-frontal projection of alpha oscillations (14, 20, 21, 31-35) to optimize network-level modulation. Intracranial current flow estimation (based on both a standard and a realistic head model) confirmed focal electrical fields in the targeted region. Simultaneously recorded EEG further confirmed alpha power increase in the targeted region as well as P→F connectivity increase at both the surface and source levels. Importantly, our mediation analysis suggests that enhancement in the alpha P→F connectivity contributes to DMN connectivity upregulation.

tACS did not affect horizontal or F→P alpha connectivity on either surface or source level (*SI Appendix*), suggesting that tACS primarily serves to amplify the underlying endogenous activity (i.e., posterior-to-frontal alpha projection). Relatedly, despite the midline montage, the effects of α-tACS on alpha posterior power and P→F connectivity emerged in the right hemisphere primarily, consistent with our previous α-tACS study using the same tACS parameters (27). Given the right-hemisphere dominance of alpha oscillations in various basic processes (41), we suspect that this hemispheric asymmetry of tACS effects is also aligned with the underlying alpha functioning. Control analyses also explored potential tACS effects outside the alpha frequency and the DMN. Analyses across the frequency spectrum (1-30 Hz) ascertained that the tACS effects were specific to the alpha frequency (*SI Appendix*). Similarly, whole-brain connectivity analysis seeded in the PCC or the tACS site further confirmed increased connectivity within the DMN and not outside the DMN (even at a lenient threshold of *p* < .01 uncorrected). These results highlight the critical contribution of alpha oscillations to the integrity of the DMN.

Increased BOLD and alpha-frequency connectivity between the tACS site and DMN hubs (PCC and mPFC; *SI Appendix*) suggest that superficial transcranial stimulation can reach the two hubs of the DMN via the tACS site and synchronize their activity, thereby inducing the upregulation of the network. Specifically, our source-based alpha connectivity analyses revealed that via the tACS site, this alpha stimulation can effectively drive alpha oscillations in the mPFC, a deep and distal structure that is critically implicated in emotional disorders but is often difficult to access via noninvasive stimulation (42). Our findings thus have important clinical implications by identifying a viable pathway for non-invasive intervention to modulate this pivotal structure.

The impact of α-tACS on DMN BOLD connectivity also reached the lateral connection between the vPCC and the rANG, suggesting that enhanced BOLD coupling of DMN midline hubs would propagate to the rest of the network. This finding could highlight the robust self-organization of an intrinsic connectivity network, enabling network-wide modulation. The anatomy of the DMN has been increasingly refined since its initial characterization. Within the “midline core”, there are actually two interdigitated subnetworks, where the PCC hub is divided into a ventral division (vPCC) and a dorsal division (dPCC), with the former connected with the mPFC hub (the anterior and ventral mPFC) and the latter with the dorsal mPFC (8, 43, 44). Accordingly, the vPCC is thought to be a primary hub for the DMN, relative to the dPCC that is likely a connector hub between the DMN and other networks (e.g., the salience network, the visual network) (44). In keeping with that, our separate examination of the ventral and dorsal subdivisions of the PCC (i.e., vPCC and dPCC) isolated tACS effects on vPCC-but not dPCC-related DMN connectivity. In fact, exploratory analysis of dPCC-seeded whole-brain connectivity revealed no clear effects of tACS (*p* < .01 uncorrected; *SI Appendix;* Fig. S2). These findings further accentuate the close association between alpha oscillations and the core of the DMN.

While mounting evidence indicates the physiological coupling and functional similarity between alpha oscillations and the DMN, they have been correlational in nature (1, 12, 15-19, 22, 23, 25). Our experimental manipulation of alpha oscillations via tACS lends mechanistic support for the DMN-alpha association, reinforcing the hypothesis that the intrinsic connectivity networks have an electrophysiological origin (4) and, specifically, that alpha oscillations underpin the DMN (14). Yet to be revealed are the precise physiological processes that underlie this association. It is possible that neural synchrony via alpha oscillations directly synchronizes BOLD fluctuations across the DMN, thereby enhancing DMN connectivity. However, as indicated by the mediation analysis (Fig. 3*B*), the effect of tACS modulation on DMN connectivity enhancement remained significant after controlling for the contribution of alpha P→F connectivity, suggesting additional mechanisms were at play.

We surmise that the DMN and alpha oscillations, two primary architectures of the brain’s organization, may be united and synergized by a more fundamental organizational architecture— the thalamocortical circuitry (45). The thalamocortical circuitry is long known to generate, sustain, and regulate alpha oscillations (46, 47). Recently, the thalamocortical circuitry has been proposed to be a key controller of the DMN (8). Particularly, thalamocortical circuits (including the thalamus, sensory cortex, and thalamic reticular nucleus) that regulate alpha oscillations are also posited to regulate the DMN by inhibiting sensory feedforward projection, regulating wakefulness and arousal, and orchestrating large-scale brain activities (8). In line with this thalamocortical hypothesis, modulation of alpha oscillations can affect the thalamocortical circuitry, which then upregulates the DMN. Future research into the interrelations between thalamocortical circuitry and the DMN, specifically through alpha activity, would further elucidate this mechanism.

That the thalamocortical circuitry could serve as the backbone of the brain’s organizational architecture (45) and regulate both the DMN and alpha oscillations would account for the prevalent thalamocortical dysfunctions, in both connectivity (thalamocortical dysconnectivity) and neural synchrony (thalamocortical dysrhythmia), in major neuropsychiatric disorders (47-49). Accordingly, therapies effectively regulating this circuitry would bring forward major breakthroughs in neuropsychiatric treatment, and non-invasive neuromodulation of alpha oscillations could be a prime candidate for such interventions.

## Materials and Methods

### Participants

41 healthy volunteers (24 female, 20.8 ± 3.2 years of age) participated in the study after providing written, informed consent approved by the Florida State University Institutional Review Board. No participants reported a history of neurological or psychiatric disorders, or current use of psychotropic medication. Participants were randomly assigned to two groups, an Active (*n* = 21) and a Sham group (*n* = 20). 2 participants (Active *n* = 2) terminated their participation prematurely due to discomfort in the scanner. 3 participants (Active *n* = 2, Sham *n* = 1) were excluded from fMRI analyses due to excessive motion (defined by > 5% of scans exceeding a frame-wise displacement index of 0.5 mm (50)), resulting in a final sample of 36 participants for fMRI analyses (Active *n* = 17, Sham *n* = 19). 4 separate participants (Active *n =* 1, Sham *n =* 3) were excluded from EEG analyses due to significant artifacts, resulting in a final sample of 35 participants for EEG analyses (Active *n* = 18, Sham *n* = 17). Participants in the two groups did not differ in age or gender distribution (*p*’s > 0.50).

### Experimental Design

The experiment consisted of three phases: Pre-stimulation RS recordings, tACS/sham stimulation, and Post-stimulation RS recordings. In both Pre- and Post-stimulation phases, participants underwent two successive 5-minute simultaneous EEG-fMRI scans (with eyes open and fixated on a central crosshair). The MR-compatible stimulation was fully integrated with simultaneous EEG-fMRI recordings such that it did not require transition between the RS recording phases.

### tACS

Alpha-frequency stimulation was administered with a ±2 mA sinusoidal current oscillating at 10 Hz using an MR-compatible High-Definition (HD) tACS system (Soterix Medical New York, NY, USA). Stimulation electrodes were placed in a 4 × 1 montage over midline occipitoparietal sites (with 4 surrounding + 1 central electrodes forming a closed circuit; Fig. 1*A*), which were selected to maximally target the primary cortical source of alpha oscillations—occipitoparietal cortex (12, 13, 27).

α-tACS or sham stimulation was administered for 20 minutes. To minimize awareness of experimental condition, the Sham group received α-tACS for 10 s at the beginning and the end of the phase. All participants completed a standard continuous performance task, which, by maintaining alertness, would enhance α-tACS efficacy (51). All participants were first told they would receive electrical stimulation and were informed of their true assignment during the debriefing at the end of experiment. Participants’ blindness to the group assignment was confirmed via a funnel interview at the debriefing, which was further corroborated with the Adverse Effects Questionnaire at the end of experiment (52). Specifically, the Active and Sham groups showed no difference in their subjective experiences during the stimulation period (*t* = 0.81, *p* = 0.423) or the degree to which they attributed these sensations to the stimulation (*t* = 1.23, *p* = 0.225).

### EEG acquisition and analyses

EEG data were recorded simultaneously with fMRI using a 64-channel MR-compatible EEG system (Brain Products GmbH, Germany). An additional electrode was placed on the participant’s upper back to record electrocardiogram (ECG) for cardioballistic artifact correction. The EEG recording system was synchronized with the fMRI scanner’s internal clock throughout acquisition to facilitate successful removal of MR gradient artifact (15). Cardioballistic and gradient artifact corrections were performed offline using an average artifact template subtraction method as implemented in Brain Vision Analyzer 2.0 (Brain Products GmbH). Details regarding EEG artifact correction and additional preprocessing are provided in the *SI Appendix*.

Power of alpha-frequency oscillations were computed using the multitaper spectral estimation technique. Directed alpha-frequency connectivity was assessed using Granger causality (GC) analysis. Source-level analysis of alpha activity was performed using the Fieldtrip toolbox implemented in SPM12 (http://www.fil.ion.ucl.ac.uk/spm/software/spm12), with the head model defined by each participant’s T1 scan. To maintain the temporal resolution needed to compute Granger causality, source-based alpha-frequency connectivity was assessed using ROI time-series derived from Exact Low Resolution Electromagnetic Tomography (eLORETA) (53). These ROI time-series were then submitted to GC analysis using the same criteria as surface-level GC analysis. Additional details are provided in the *SI Appendix*.

### MRI Acquisition and Preprocessing

Gradient-echo T2-weighted echoplanar images were acquired on a 3T Siemens Prisma MRI scanner using a 64-channel head coil with axial acquisition. Imaging parameters included TR/TE: 1800/22.40 ms; slice thickness 1.8 mm, gap .45 mm; and in-plane resolution/voxel size 1.8 × 1.8 mm, multiband acceleration factor = 2 (54); GRAPPA acceleration factor = 2. A high-resolution (.9 × .9 × .9 mm3) 3D-MPRAGE T1 scan and an echoplanar field map were also required. Imaging data were preprocessed using SPM12, including slice-time correction, spatial realignment, fieldmap correction, and normalization using Diffeomorphic Anatomical Registration Through Exponentiated Lie algebra (DARTEL). To further remove artifacts potentially contributing to spurious RS activity variance (50), we implemented additional preprocessing using the DPARSFA toolbox: (1) mean centering and whitening of timeseries; (2) temporal bandpass (.01-.08 Hz) filtering; (3) general linear modeling (GLM) to partial out head motion with 24 nuisance variables (six head motion parameters each from the current and previous scan and their squared values); and (4) scrubbing of significant motion (“spikes”) based on framewise displacement index (FDi > .5 mm). Further details are provided in the *SI Appendix*.

### Regions of Interest (ROIs)

Regions of interest (ROIs) consisted of key nodes of the DMN, including the midline hubs, posterior cingulate cortex (PCC) and medial prefrontal cortex (mPFC), and the bilateral angular gyri (ANG) (55). Masks for the mPFC and ANG ROIs were drawn from the Willard Atlas (56). The PCC hub consists of functionally-dissociable ventral and dorsal subdivisions; specifically, the ventral PCC (vPCC) primarily linking DMN nodes and the dorsal PCC (dPCC) with extensive extra-DMN connections (8, 44). We thus extracted masks for the vPCC and dPCC sub-divisions individually from the Brainnetome Atlas (57). The site of tACS was also included as an ROI, represented by a 10-mm sphere centered on the voxel with the maximal electric field (x, y, z = 16, -86, 36; Fig. 1*A*).

### Resting-state functional connectivity (rsFC)

Pre- and Post-stimulation fMRI timeseries from each of the six ROIs were submitted to Pearson’s correlation analysis to construct a 5×5 correlation matrix for each session for each participant. The pair-wise correlation coefficients were Fisher Z-transformed before submission to statistical analyses. Seed-based whole-brain connectivity, with PCC and tACS site as seeds, was further evaluated to ascertain the extent of connectivity changes.

### Statistical analysis

We first established the efficacy of tACS by examining changes in alpha power and P→F alpha connectivity. We then conducted ROI-based connectivity analyses for fMRI rsFC. These effects of tACS were evaluated with simple contrasts (paired t-tests of Pre vs. Post sessions) in the Active group (*p* < .05). To control for time-related confounds, we also performed double contrasts of Pre versus Post between Active and Sham groups (Active – Sham_Post – Pre_; *p* < .05). Given the multiple ROIs considered in the analyses, we applied false discovery rate/FDR correction to the double contrasts across the ROIs. Seed-based whole-brain connectivity was corrected with small volume correction/SVC (FDR *p* < .05) in SPM12. For ROIs that emerged from the rsFC analysis, we submitted their source-level alpha frequency GC to the same simple and double contrasts. To link EEG and fMRI effects of tACS, we further submitted them into Pearson correlation analyses (*p* < .05). Significant EEG-fMRI correlations were followed by mediation analyses, to elucidate the contribution of alpha enhancement to α-tACS effects on DMN connectivity. The PROCESS macro for SPSS (58) was used to estimate 5,000 bias-corrected bootstrap samples, from which a 95% confidence interval (CI) was created to test the indirect effect of α-tACS on DMN connectivity through alpha activity.

## Acknowledgments

This research was supported by the National Institute of Mental Health grants R01MH093413 (W.L.) and the FSU Chemical Senses Training (CTP) Grant Award T32DC000044 (K.C.) from the National Institutes of Health (NIH/NIDCD), the National High Magnetic Field Laboratory (J.T.R.) which is supported by the National Science Foundation through NSF (DMR-1644779 and *DMR-1157490)* and the State of Florida.).

