## Supplementary Information for "Transcranial stimulation of alpha oscillations upregulates the default mode network"

**This PDF file includes:**

Supplementary text
Figures S1 and S2
SI References

### Supplementary Information Text

### Supplementary Methods

#### tACS current distribution in a realistic head model

Individual differences in head anatomy have been demonstrated to affect the current distribution of transcranial stimulation. To validate our estimated current flow from a standard head model, a finite-element model simulation was performed using the average T1 structural image from the Active group through the Realistic vOlumetric-Approach to Simulate Transcranial Electric Stimulation (ROAST) pipeline (1). The model consisted of stimulation electrodes placed at P1, P2, O1, and O2 with a net current of .5 mA each, and a central POz electrode with a net current of 2 mA. Default tissue conductance parameters and stimulation current parameters were used. The results from the individualized head model demonstrated highly overlapping current distributions with the standard head model (peak coordinate: x, y, z = -16, -89, 33; Fig. S1).

#### EEG Preprocessing

Cardioballistic and gradient artifact corrections were performed offline using an average artifact template subtraction method as implemented in Brain Vision Analyzer 2.0 (Brain Products GmbH). The gradient artifact template was constructed with a sliding-window approach over 41 consecutive volumes. For cardioballistic artifact correction, ECG R peaks were first identified using a semi-automatic detection approach (i.e., automated peak detection combined with visual inspection) and then used to construct a delayed average artifact template over 41 consecutive heartbeat events, which was then subtracted from the EEG data. Artifact-corrected EEG data were then band-pass filtered between 0.5 and 50 Hz and down-sampled to 250 Hz before submission to the *Fully Automated Statistical Thresholding for EEG artifact Rejection* (FASTER) algorithm for further artifact correction (2). The FASTER algorithm (implemented in EEGLAB) corrected EEG data for remaining physiological and non-physiological artifacts in each channel, epoch, and independent component. Output data were epoched into segments of 1.8 s (i.e., TR length), centered on the onset of each fMRI scan.

*EEG Power Analyses*

Power of alpha-frequency oscillations was computed for each electrode within each epoch using the multitaper spectral estimation technique (3). Alpha power was normalized by the mean power of the entire frequency spectrum (1-50 Hz) within each epoch, and then averaged across occipitoparietal electrodes where alpha is maximally distributed (Fig. 1c).

*EEG directed alpha-frequency connectivity (Granger causality)*

Directed alpha-frequency connectivity was assessed using Granger causality (GC) analysis (4, 5). EEG data were transformed to reference-free current source density data (CSD) using the surface Laplacian algorithm (6). CSD data from ipsilateral posterior-frontal pairs (7) were then submitted to bivariate autoregressive (AR) modeling, from which Granger causality spectra were derived (4) and averaged across the alpha frequency (8-12 Hz) for each posterior-frontal pairs. A model order of 20 (80 msec in time for a sampling rate of 250 Hz) was chosen in a two-step process: 1) Akaike Information Criteria (AIC) and 2) comparing spectral estimates obtained from the Fourier-based AR model on data pooled across all subjects (8). Given *a priori* hypotheses on the posterior→frontal dominance of alpha-frequency connectivity at rest (7, 9-11) and the posterior target of the α-tACS, analyses were constrained to the posterior→frontal direction.

*EEG source analysis*

Source-level analysis of alpha activity was performed using the Fieldtrip toolbox implemented in SPM12 (http://www.fil.ion.ucl.ac.uk/spm/software/spm12), with the head model defined by each participant's T1 scan (12). The multiple sparse priors algorithm was used to generate the inverse solution. Induced alpha power was extracted using a series of Morlet wavelet projectors within each individual epoch and then averaged across epochs, as implemented by default in SPM12.

To maintain the temporal resolution needed to compute Granger causality, source-based alpha-frequency connectivity was assessed using ROI time-series derived from Exact Low Resolution Electromagnetic Tomography (eLORETA) (13). These ROI time-series were then submitted to Granger causality analysis using the same criteria described above. ROIs consisted of 10-mm spheres around bilateral cortical DMN hubs – posterior cingulate cortex (PCC), medial prefrontal cortex (mPFC) and angular gyrus (ANG) – and the maximal tACS stimulation site.

**MRI Acquisition and Preprocessing**

Gradient-echo T2-weighted echoplanar images (348 scans) were acquired on a 3T Siemens Prisma MRI scanner using a 64-channel head coil with axial acquisition. Imaging parameters were TR/TE: 1800/22.40 ms; flip angle: 40°, field of view 212 mm, slice thickness 1.8 mm, gap .45 mm; in-plane resolution/voxel size 1.8 × 1.8 mm; multiband acceleration factor = 2; GRAPPA acceleration factor = 2 (14-16). A high-resolution (.9 × .9 × .9 mm3) 3D-MPRAGE T1 scan and an echoplanar field map were also required. Imaging data were preprocessed using SPM12 and the DPARSFA toolbox (17), including the removal of the first 12 dummy scans, slice-time correction, spatial realignment, and normalization using Diffeomorphic Anatomical Registration Through Exponentiated Lie algebra (DARTEL; (18)). To further remove artifacts potentially contributing to spurious RS activity variance (19, 20), we implemented additional preprocessing: (1) mean centering and whitening of timeseries; (2) temporal bandpass (.01-.08 Hz) filtering; (3) general linear modeling (GLM) to partial out head motion with 24 nuisance variables (six head motion parameters each from the current and previous scan and their squared values); and (4) scrubbing of significant motion (“spikes”) based on framewise displacement index (FDi) [FDi= |Δdix| + |Δdiy| + |Δdiz| + |Δαi| + |Δβi| + |Δγi|]; Scans with FDi over 0.5 mm were classified as spikes and removed (19).

**Supplementary Results**

**Specificity of α-tACS to the alpha frequency and P→F connectivity.** To ascertain the specificity of α-tACS effects to the alpha frequency, we evaluated Post-Pre contrasts of EEG power in the Active group for neighboring frequency bins. No effects emerged in the delta (1-3 Hz; $t$ = -0.970, $p$ = 0.345), theta (4-7 Hz; $t$ = -0.864, $p$ = 0.400), or beta (12-30 Hz; $t$ = -1.476, $p$ = 0.158) frequencies. A similar contrast was performed for P→F connectivity, with again no effects emerging in delta ($t$ = -0.176, $p$ = 0.862), theta ($t$ = 0.479, $p$ = 0.638), or beta ($t$ = 0.756, $p$ = 0.460) frequencies.

To further ascertain the specificity of the P→F effects, similar contrasts of Post-Pre alpha-frequency F→P directed connectivity were performed in the Active group. No effects emerged for F→P connectivity in the left ($t$ = -0.769, $p$ = 0.848) or right ($t$ = -0.151, $p$ = 0.882) hemisphere. No effects emerged for F→P connectivity in neighboring frequencies, as well ($p$’s > 0.351).

Finally, to demonstrate the unique association between alpha connectivity and DMN connectivity, Pearson correlation analyses were performed between Post-Pre changes in DMN connectivity and P→F connectivity in neighboring frequencies. No associations emerged between changes in vPCC-mPFC connectivity and delta ($r$ = 0.129, $p$ = 0.482), theta ($r$ = 0.249, $p$ = 0.170), or beta ($r$ = 0.288, $p$ = 0.110) P→F connectivity. A trending association between increases in vPCC-rANG connectivity and decreases in beta P→F connectivity emerged ($r$ = -0.333, $p$ = 0.062). No association emerged with the delta ($r$ = 0.083, $p$ = 0.652) or theta ($r$ = 0.028, $p$ = 0.878) frequencies.

**tACS effects on whole-brain PCC-seed-based connectivity.** Whole-brain vPCC seed-based connectivity maps revealed similar results (Figure 2C). A double contrast of Time (post- minus pre-stimulation) and Group (active minus sham) on whole-brain vPCC seed connectivity revealed bilateral clusters in the mPFC: left mPFC (k = 13, peak voxel MNI coordinates: -12, 46, 0; $Z_{34}$ = 3.64, $p$ < 0.001, SVC $p_{FWE}$ = 0.103); right mPFC (k = 17, peak voxel MNI coordinates: 8, 46, 0; $Z_{34}$ = 4.48, $p$ < 0.001, SVC $p_{FWE}$ = 0.063; Figure 2C), as well as a cluster within the rANG (k = 25, peak voxel MNI coordinates: 52, -50, 24; $Z_{34}$ = 3.94, $p$ < 0.001, SVC $p_{FWE}$ = 0.030). Follow-up contrasts of Time for each Group revealed active-specific increases in vPCC connectivity with the left mPFC (k = 29, peak voxel MNI coordinates: -12, 46, 0; $Z_{16}$ = 4.07, p < 0.001, SVC $p_{\mathrm{FWE}}$ = 0.021) and right mPFC (k = 16, peak voxel MNI coordinates: 4, 46, -4; $Z_{16}$ = 3.76, p < 0.001, SVC

$p_{FWE}$ = 0.076), as well as the rANG (k = 15, peak voxel MNI coordinates: 54, -50, 24; $Z_{16}$ = 4.12, $p$ < 0.001, SVC $p_{FWE}$ = 0.014). No such increases emerged in the sham group (Z's < 1.68, SVC $p_{FWE}$ > 0.997).

Whole-brain dPCC seed-based connectivity maps were further evaluated to ascertain potential changes in connectivity with non-DMN nodes, as the dPCC demonstrates greater inter-network connectivity compared to the vPCC. Consistent with this notion, whole-brain dPCC seed-based connectivity maps revealed no effects in our *a priori* DMN ROIs, as well as no substantial clusters in hubs of other resting state networks, even at a more liberal threshold for visualization ($p$ < 0.01 uncorrected, $k$ > 10; Fig. S2).

**tACS effects on whole-brain $tACS_{site}$-seed-based connectivity**. Whole-brain $tACS_{site}$ seed-based connectivity maps revealed similar results (Fig. 4*B*). A double contrast (Post - $Pre_{Active - Sham}$) revealed clusters within the mPFC (k = 5, peak voxel MNI coordinates: -2, 46, 8, $Z_{34}$ = 3.72, $p$ < 0.001, SVC $p_{FWE}$ = 0.078) and vPCC (k = 9, peak voxel MNI coordinates: 2, -52, 20, $Z_{34}$ = 3.10, $p$ < 0.001, SVC $p_{FWE}$ = 0.277). Follow-up contrasts of Time for each Group revealed Active-specific increases in $tACS_{site}$ connectivity with the right mPFC (k = 23, peak voxel MNI coordinates: -10, 48, -2; $Z_{16}$ = 3.40, p < 0.001, SVC $p_{FWE}$ = 0.243) and right vPCC (k = 2, peak voxel MNI coordinates: 2, -52, 30; $Z_{16}$ = 3.72, p < 0.001, SVC $p_{FWE}$ = 0.051). No effects emerged in the right angular gyrus. No such increases were seen in the Sham group (Z's < 1.68, SVC $p_{FWE}$ > 0.997).

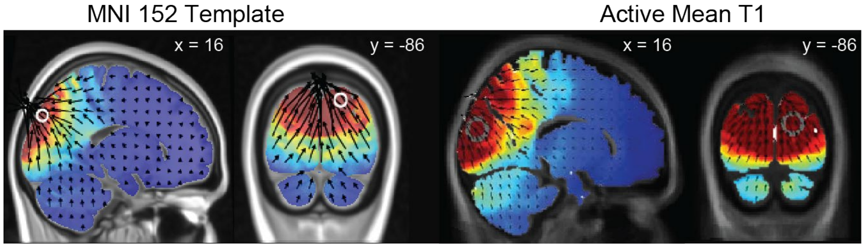


**Fig. S1. Standard versus realistic head model.** Finite-element estimation of the current flow based on a standard head model (Left) versus the average T1 from the Active group (Right) demonstrated a high degree of concordance.

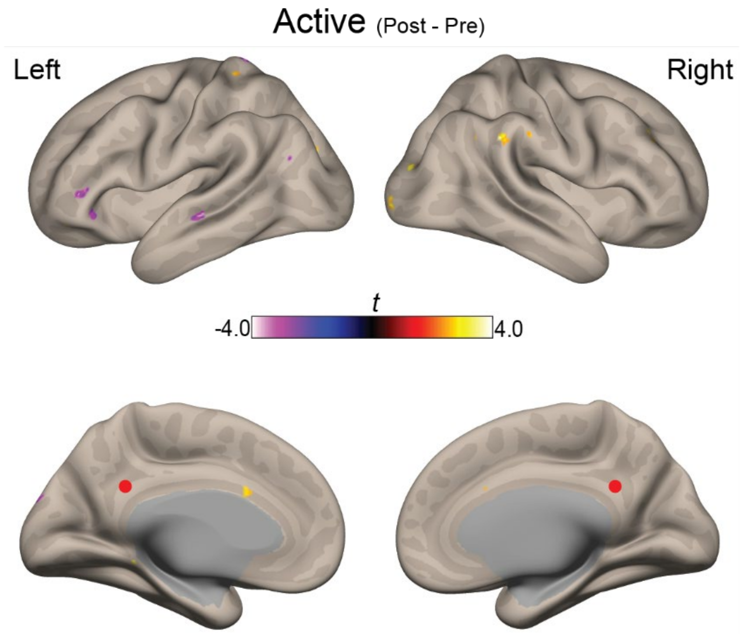


**Fig. S2. No effect of α-tACS on dPCC connectivity.** dPCC-seed whole-brain connectivity maps indicate limited changes in connectivity with other RSNs, even at a liberal threshold ($p < 0.01$ uncorrected, $k > 10$). Red dot indicates dPCC seed.
